# Divergent effects of healthy ageing on semantic knowledge and control: Evidence from novel comparisons with semantically-impaired patients

**DOI:** 10.1101/216192

**Authors:** Paul Hoffman

## Abstract

Effective use of semantic knowledge requires a set of conceptual representations and control processes which ensure that currently-relevant aspects of this knowledge are retrieved and selected. It is well-established that levels of semantic knowledge increase across the lifespan. However, the effects of ageing on semantic control processes have not been assessed. I addressed this issue by comparing the performance profiles of young and older people on a verbal comprehension test. Two sets of variables were used to predict accuracy and RT in each group: (a) the psycholinguistic properties of words probed in each trial, (b) the performance on each trial by two groups of semantically-impaired neuropsychological patients. Young people demonstrated poor performance for low frequency and abstract words, suggesting that they had difficulty processing words with intrinsically weak semantic representations. Indeed, performance in this group was strongly predicted by the performance of patients with semantic dementia, who suffer from degradation of semantic knowledge. In contrast, older adults performed poorly on trials where the target semantic relationship was weak and distractor relationships strong – conditions which require high levels of controlled processing. Their performance was not predicted by the performance of semantic dementia patients, but was predicted by the performance of patients with semantic control deficits. These findings indicate that the effects of ageing on semantic cognition are more complex than has previously been assumed. While older people have larger stores of knowledge than young people, they appear to be less skilled at exercising control over the activation of this knowledge.

## Introduction

Semantic knowledge, for the meanings of words and properties of objects, shapes our understanding of the environment and guides our behaviour. In addition to storing this information in memory, effective semantic cognition – that is, the ability to use semantic knowledge to complete cognitive tasks – requires us to *control* how we retrieve and use it in specific situations (Badre & Wagner, 2002; Hoffman, McClelland, & Lambon Ralph, 2018; Jefferies, 2013; Lambon Ralph, Jefferies, Patterson, & Rogers, 2017; Yee & Thompson-Schill, 2016). Cognitive control over the activation and selection of semantic information is critical because we store a wide range of information about any particular concept and different aspects of this knowledge are relevant in different circumstances (Hoffman, Lambon Ralph, & Rogers, 2013; Hoffman et al., 2018; Saffran, 2000; Thompson-Schill, D’Esposito, Aguirre, & Farah, 1997). If a chef is searching her kitchen for a lemon, for example, knowledge about its colour will be helpful in guiding her behaviour. However, when she begins to cook with the lemon, colour is less critical and she must instead retrieve information about its scent and flavour. This ability to retrieve task-relevant semantic information while avoiding interference from competing elements of knowledge is often termed “semantic control”.

How do semantic abilities change across the lifespan? There is good evidence that the amount of semantic knowledge people have, as indexed by their scores on vocabulary tests, increases as they grow older and remains relatively stable into old age (Grady, 2012; Nilsson, 2003; Nyberg, Bäckman, Erngrund, Olofsson, & Nilsson, 1996; Park et al., 2002; Rönnlund, Nyberg, Bäckman, & Nilsson, 2005; Salthouse, 2004). A meta-analysis of 210 studies indicated that adults aged over 60 score substantially higher on vocabulary tests than young adults (Verhaeghen, 2003). This effect is typically explained in terms of older participants continuing to add to their semantic knowledge store throughout their lives. Consistent with this conclusion, greater time spent in formal education appears to account for much of the age effect (Verhaeghen, 2003).

However, while it seems clear that older people have a larger repository of semantic knowledge than young people, the effect of ageing on semantic control processes has rarely been investigated. Other forms of cognitive control, such as response inhibition, task-switching and prevention of memory intrusions, tend to deteriorate as people grow older (Borella, Carretti, & De Beni, 2008; De Beni & Palladino, 2004; Hasher & Zacks, 1988; Salthouse & Miles, 2002; Treitz, Heyder, & Daum, 2007; Verhaeghen & Cerella, 2002) and it is possible that the cognitive control of semantic knowledge suffers a similar fate. This remains a largely unexplored possibility in the cognitive ageing literature. The relationship between semantic control and cognitive control in other domains is complex, with researchers proposing that some aspects of the regulation of semantic knowledge are performed by domain-general systems for competition resolution while others require more specialised neural resources (Badre, Poldrack, Pare-Blagoev, Insler, & Wagner, 2005; Jefferies, 2013; Nagel, Schumacher, Goebel, & D‘Esposito, 2008; Whitney, Kirk, o’Sullivan, Lambon Ralph, & Jefferies, 2011). The relationship between semantic and non-semantic control was not the focus of the present investigation, but it is important to note that the degree to which these processes share neural resources is likely to determine the extent to which they decline in parallel in ageing.

Although studies of cognitive ageing have not investigated semantic control processes in detail, there has been a stronger focus on these abilities in the neuropsychological literature. A number of studies have indicated that semantic knowledge representations and the semantic control system can be impaired independently following brain damage (Jefferies & Lambon Ralph, 2006; Rogers, Patterson, Jefferies, & Lambon Ralph, 2015; Warrington & Cipolotti, 1996). The clearest example of impairment to semantic knowledge is the syndrome of semantic dementia (SD), in which patients exhibit a progressive and highly selective loss of semantic representations (Patterson, Nestor, & Rogers, 2007). The erosion of semantic knowledge in this condition follows a predictable pattern whereby patients are more likely to retain information that is experienced more frequently, is more typical of its domain and is shared across many different concepts (Funnell, 1995; Hoffman, Jones, & Lambon Ralph, 2012; Mayberry, Sage, & Lambon Ralph, 2011; Rogers et al., 2015; Woollams, Cooper-Pye, Hodges, & Patterson, 2008). Such information is considered to be represented more strongly in the semantic store (Rogers et al., 2004; Rogers & McClelland, 2004). Some studies have also found that SD patients have poorer knowledge for abstract words, relative to concrete ones (Hoffman & Lambon Ralph, 2011; Jefferies, Patterson, Jones, & Lambon Ralph, 2009), which has been attributed to concrete words benefiting from a richer set of sensory-motor associations (Hoffman, 2016). The loss of semantic knowledge in this condition appears to be a direct consequence of atrophy to the anterior temporal cortices (Butler, Brambati, Miller, & Gorno-Tempini, 2009; Mion et al., 2010). Studies using functional neuroimaging and brain stimulation in healthy individuals also implicate this region in the representation of semantic knowledge (Humphreys, Hoffman, Visser, Binney, & Lambon Ralph, 2015; Pobric, Jefferies, & Lambon Ralph, 2007; Rogers et al., 2006; Visser, Jefferies, & Lambon Ralph, 2010).

The deterioration of semantic knowledge in SD has been contrasted with multimodal semantic deficits that can occur following left-hemisphere stroke (Crutch & Warrington, 2008; Jefferies & Lambon Ralph, 2006; Rogers et al., 2015; Warrington & Cipolotti, 1996). Stroke is less likely to affect the anterior temporal cortices, which benefit from a double arterial blood supply (Gloor, 1997). Instead, semantic deficits in this group of patients, who are often termed semantic aphasics (SA), result from prefrontal and posterior temporoparietal damage (Noonan, Jefferies, Corbett, & Lambon Ralph, 2010). Consistent with this different locus of damage, semantic knowledge representations appear to be intact in SA patients. Instead, their semantic deficits stem from difficulty in controlling how this knowledge is accessed and selected according to current task demands. For example, they have difficulty identifying meaningful relationships between words where they do not share a strong automatic association (e.g., they can detect the semantic relationship between *necklace* and *earring* but not between *necklace* and *trousers*) (Noonan et al., 2010). They also perform poorly when required to ignore strong, pre-potent semantic associations that are irrelevant to the task at hand. For example, in a synonym matching task they will erroneously select antonyms if they have a stronger relationship with the probe (e.g., matching *major* with *minor* rather than *important*). Thus, SA patients find semantic tasks particularly challenging when the target semantic relationship is weak and any irrelevant non-target relationships are strong (Jefferies & Lambon Ralph, 2006). Functional neuroimaging studies indicate that these conditions also elicit the strongest activation in prefrontal and temporoparietal regions associated with control of semantic processing (Badre et al., 2005; Noonan, Jefferies, Visser, & Lambon Ralph, 2013; Thompson-Schill et al., 1997).

Like SD patients, individuals with SA find it easier to understand highly concrete words (Hoffman, Jefferies, & Lambon Ralph, 2010) but, unlike SD patients, they show little or no effects of frequency in their comprehension (Almaghyuli, Thompson, Lambon Ralph, & Jefferies, 2012; Hoffman, Jefferies, & Lambon Ralph, 2011a; Hoffman, Rogers, & Lambon Ralph, 2011b; Warrington & Cipolotti, 1996). Instead, they show particularly poor comprehension of words that are semantically diverse, i.e., words that can be used in a wide variety of different contexts (Hoffman et al., 2011b). The contextual promiscuity of these words appears to pose problems for SA patients because they have difficulty selecting which aspect of the word’s meaning is relevant for the current context (see also Noonan et al., 2010). In contrast, semantic diversity has no effect on comprehension in SD patients (Hoffman et al., 2011b).

Comparative studies of SD and SA patients have provided us with a clear model of the characteristics of semantic processing that are associated with weakness in either the representation of semantic knowledge or in its controlled processing. The present study used these insights to investigate areas of semantic weakness in healthy young and older adults. Participants completed a forced-choice verbal comprehension task in which they were asked to identify the synonyms of 96 words. The aim of the study was to investigate which factors influenced performance in each group. As summarised above, neuropsychological deficits in semantic knowledge and control are associated with sensitivity to different psycholinguistic variables. Accordingly, we first investigated which psycholinguistic properties best predicted performance in young and older people. We predicted that young people, like SD patients, would show larger effects of word frequency, as they are generally assumed to have a less developed store of semantic representations. Conversely, if older people have difficulties in the control of semantic processing, we would expect the factors influencing their performance to be similar to those that predict comprehension in SA: i.e., the semantic diversity of the words probed and the relative strength of target and non-target semantic relationships.

As a second test of these hypotheses, we made use of previously-published data from 26 SD and SA patients who completed the same comprehension task. For each trial in the task, we computed an SD performance index and an SA performance index, which indicated how likely patients from either group were to respond correctly to the stimulus. These provided a measure of how difficult each trial was for an individual with impairment of semantic representation and for someone with an impairment of semantic control. We then used these measures to predict performance in our healthy participants. We expected that both of these indices would predict performance; in other words, we expected healthy individuals to respond more quickly and accurately on the trials that the patients tended to do well on. Importantly, however, we predicted that the two indices would have different effects in the two age groups. We hypothesised that the SD performance index would be a stronger predictor of performance in young people, since this index should measure representational difficulty and young people are assumed to have less-developed semantic representations. Conversely, if older people are less efficient in exercising semantic control, we would expect the SA performance index to be the stronger predictor of performance, since this measure would act as a marker of the semantic control demands.

## Method

### Participants

Twenty-seven young adults, aged between 18 and 22, were recruited from an undergraduate Psychology course and participated in the study in exchange for course credit. Twenty-seven older adults, aged between 61 and 90, were recruited from the Psychology department’s volunteer panel. The panel consists of individuals from the [LOCATION REDACTED] area who have volunteered to take part in research studies. As they are self-selecting, such individuals are not necessarily representative of the population at large and may represent a particularly well-educated sample. Young and older adults did not differ significantly in years of education completed (Mann-Whitney *U* = 264, *p* = 0.17), however, suggesting high but equivalent levels of formal education in both groups (see Table 1). All participants reported to be in good health with no history of neurological or psychiatric illness. Experience and competence in computer use was not formally assessed and may have differed between groups. It is also important to note that age range of the older group was greater than that of the younger group.

**Table 1:** Demographic information and mean test scores for young and older participants

|  | Young adults | Older adults |
| --- | --- | --- |
| N | 27 | 25 |
| Age | 18.9 (0.8) | 74.9 (9.3)*** |
| Sex M:F | 10:17 | 11:14 |
| Years of education | 13.9 (0.8) | 13.4 (3.0) |
| MMSE /30 | 28.6 (0.9) | 29.0 (1.2)* |
| Wisconsin card-sorting task /64 | 50.8 (6.0)*** | 36.3 (11.7) |
| Trails A time (s) | 30.8 (10.7) | 36.3 (11.8) |
| Trails B time (s) | 48.7 (13.1)** | 69.0 (24.1) |
| Category fluency (items per category) | 23.1 (5.5) | 21.2 (5.0) |
| Letter fluency (items per category) | 14.4 (5.0) | 15.3 (5.5) |
| Spot the Word test /60 | 47.0 (3.8) | 54.3 (3.4)*** |
| Mill Hill test (modified) /44 | 16.6 (4.4) | 27.5 (5.5)*** |

*Standard deviations are shown in parentheses. Asterisks indicate the significance of Mann-Whitney U tests comparing young and older adults. \* = $p < 0.05$ ; \*\* = $p < 0.01$ ; \*\*\* = $p < 0.001$ .*

### General cognitive assessments

Participants completed a series of tests of general cognitive function and executive ability, to provide some basic information on the cognitive abilities of each group. The Mini-Mental State Examination was used as a general cognitive screen. Participants scoring less than 26/30 were excluded from the study. Task-switching ability was assessed using the Trail-making task (Reitan, 1992). Executive function was also assessed with a computerised version of the Wisconsin Card-Sorting Test, consisting of 64 trials (Mueller & Piper, 2014). Three categories of verbal fluency were administered, in which participants were given one minute to produce as many words as possible that fit a specific criterion. The criteria included two semantic categories (animals and household objects) and one letter of the alphabet (words beginning with F). Participants completed two standardised tests of vocabulary knowledge: the Spot-the-Word Test from the Speed and Capacity of Language Processing battery (Baddeley, Emslie, & Smith, 1992) and a modified version of the Mill Hill vocabulary test (Raven, Raven, & Court, 1989), in which participants were asked to select the synonyms of low-frequency words from four alternatives.

### Materials

Participants completed a 96-item synonym judgement test, which has been used in a number of previous studies to assess verbal comprehension in healthy and impaired populations (e.g., Almaghyuli et al., 2012; Hoffman & Lambon Ralph, 2011; Hoffman et al., 2011b; Jefferies et al., 2009). On each trial, participants are presented with a probe word and asked to select which of three alternatives is most closely related in meaning to the probe (see Figure 1 for examples). No words were repeated during the test. The test was designed to vary the frequency and imageability of the words probed in a 2 (frequency) × 3 (imageability) factorial design. In the present study, these variables were entered into model as parametric predictors of performance, along with others (see below). Probes and targets were intended to be synonyms (e.g., forest-woods) or close categorical neighbours (e.g., frog-toad). Associative relationships (e.g., dog-bone) were avoided. The full set of stimuli are provided as Supplementary Materials.

**Figure 1:**
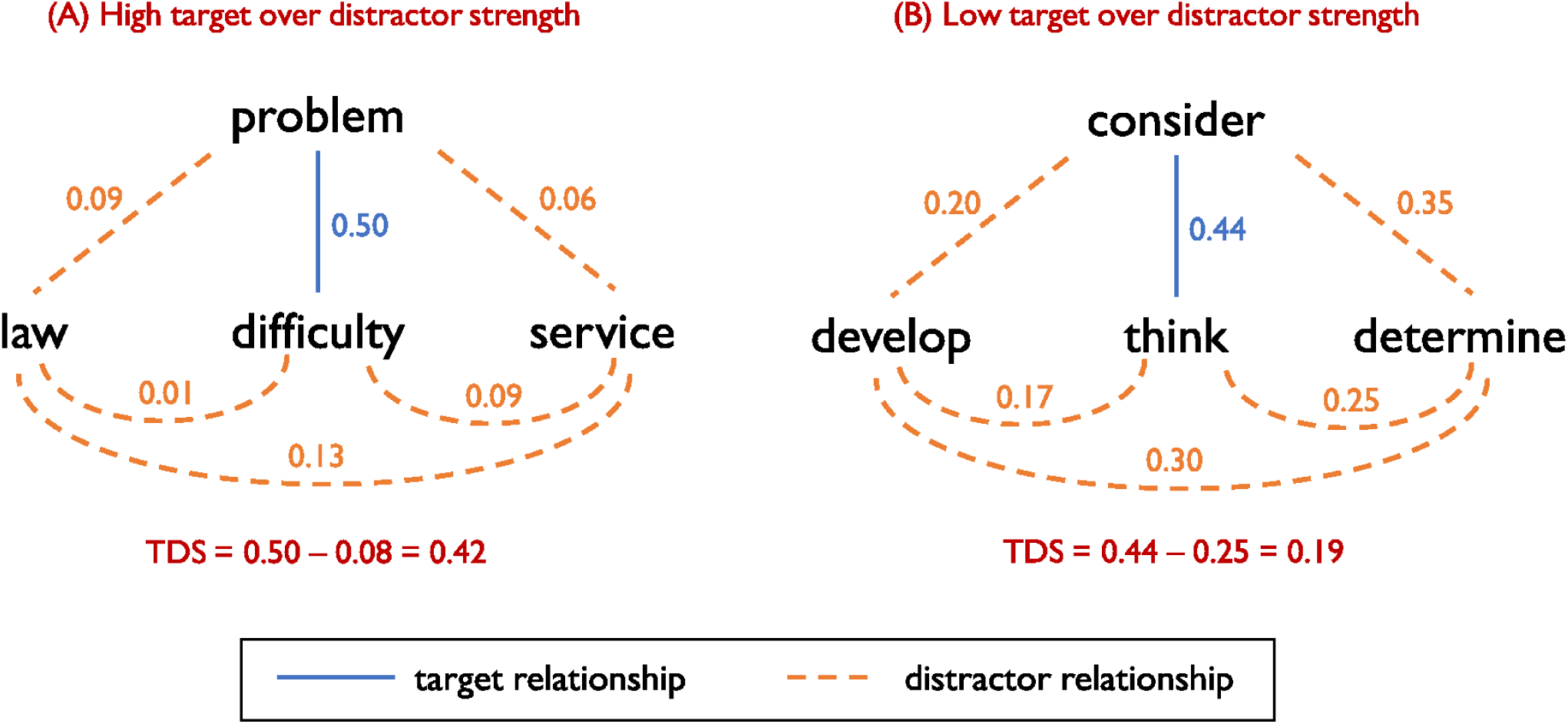
Examples of trials with high and low TDS values

### Procedure

All participants completed the background neuropsychological tests before completing the synonym judgement task. The synonym judgement task was presented on a PC running E-prime software. Each trial began with a fixation cross presented for 500ms. The probe then appeared in the centre of the screen with the three options in a line below. Participants indicated their choice by button press (1,2 or 3 for left, middle or right) and were instructed to respond as quickly as possible without making mistakes. No time limit was placed on responses. When a response was registered, the next trial began after a 500ms delay. The position of the target (left, centre, right) was randomised on each trial. The order of trials was also randomised for each participant. The main test was preceded by ten practice trials.

### Analyses

Analyses were performed on accuracy and RT data. Mann Whitney U tests were first performed to test for group differences on the task as a whole. Subsequent analyses used mixed effects models to predict accuracy and RT at the level of individual trials. The first set of analyses used the psycholinguistic properties of the trials as predictors. Separate mixed models were estimated for accuracy and RT in young and older participants. In addition, combined models including all participants were estimated. These additionally included effects of age group and the interaction of group with the other predictors. The psycholinguistic predictors were as follows:

#### Frequency

Log word frequencies for the probe items were obtained from the CELEX database (Baayen, Piepenbrock, & van Rijn, 1993).

#### Imageability

Imageability ratings for the probe items were taken from the MRC psycholinguistic database (Coltheart, 1981). These varied on a scale from 100 to 700.

#### Semantic diversity

Values for the probe items were obtained from Hoffman et al. (2013). This semantic diversity measure is based on a large corpus of natural language and indexes the degree to which a word is used in a wide range of different contexts in the corpus.

#### Target vs. distractor strength (TDS)

This measure was devised for the present study and was designed to quantify characteristics of the test that influence the need for semantic control. The measure indexed the strength of the target semantic relationship on each trial, relative to irrelevant relationships involving the distractor items. Calculation of this measure for two example trials are shown in Figure 1. Calculation of the TDS measure required quantitative measures of the strength of the semantic relationships between word pairs, which we obtained from a publically available set of word representations generated by the word2vec neural network (Mikolov, Chen, Corrado, & Dean, 2013; downloaded from https://code.google.com/archive/p/word2vec/). We will now describe this model in brief.

In common with other distributional models of word meaning including latent semantic analysis (Landauer & Dumais, 1997), the word2vec model represents words as high-dimensional vectors, where similarity in two words’ vectors indicates that they appear in similar contexts, and thus are assumed to have related meanings. The vectors were generated by training the network on the 100-billion word Google News corpus. Each time the network was presented with a word from the corpus, it was trained to predict the context in which it appeared, where context was defined as the two words preceding and following it in the corpus. As a result of this training, the model learns to represent words used in similar contexts with similar patterns. We used these representations because a recent study has shown that they outperform other available vector datasets in predicting human semantic judgements (Pereira, Gershman, Ritter, & Botvinick, 2016).

The strength of the semantic relationship between two words was defined as the cosine similarity of their word2vec vectors. This value was calculated for all pairs of words in all trials. TDS was defined as the strength of the target relationship minus the mean of all distractor relationships. As outlined in the Introduction, semantic control demands are highest when the target semantic relationship is relatively weak and there is competition from strong distractor relationships. Thus, trials with low TDS values were assumed to place the greatest demands on control processes.

The second set of analyses used previously published performance data from SD and SA patients to predict performance in healthy participants. Hoffman et al. (2011b) reported data on the 96-item synonym judgement from 13 SD and 13 SA patients. Demographic information and scores on a range of neuropsychological tests are reproduced in Table 2. As previously reported by Hoffman et al., the SD patients were recruited from memory clinics in Bath, Cambridge, Liverpool and Manchester, UK and presented with a progressive and selective multi-modal impairment of semantic memory. They fulfilled contemporary diagnostic criteria for SD (Neary et al., 1998) and imaging in each case indicated bilateral atrophy of the anterior temporal lobes. The 13 SA patients were recruited from stroke clubs and speech and language therapy services in the Manchester (UK) area. The inclusion criteria, originally described by Jefferies and Lambon Ralph (2006), were that each patient had suffered a left-hemisphere stroke at least one year previously and that they showed impairments in both picture and word versions of a semantic association test (Bozeat, Lambon Ralph, Patterson, Garrard, & Hodges, 2000). Thus, both SD and SA patients presented with multi-modal semantic deficits. Indeed, both groups of patients experience word-finding difficulties and impairments of object use which have been linked to their underlying semantic deficits (Corbett, Jefferies, Ehsan, & Lambon Ralph, 2009; Jefferies & Lambon Ralph, 2006).

**Table 2:** Demographic information and neuropsychological test scores for patients, compared with normative data from healthy individuals

|  | SA | SD | Normative data |
| --- | --- | --- | --- |
| N | 13 | 13 |  |
| Age | 66.2 (12.5) | 63.4 (7.2) |  |
| Education (leaving age) | 15.6 (1.7) | 16.0 (2.6) |  |
| *Semantic processing* |  |  |  |
| Synonym judgement task /96 | 68.7 (10.5) | 69.3 (10.8) | 94.5 (1.8) |
| Picture naming<sup>a</sup> /64 | 28.2 (21.7) | 34.3 (17.7) | 62.3 (1.6) |
| Word-picture matching<sup>a</sup> /64 | 51.2 (10.3) | 51.7 (14.4) | 63.8 (1.4) |
| Camel and Cactus test: words<sup>a</sup> /64 | 39.3 (10.2) | 42.3 (12.2) | 60.7 (2.1) |
| Camel and Cactus test: pictures<sup>a</sup> /64 | 38.2 (11.9) | 44.2 (12.1) | 59.0 (3.1) |
| Category fluency<sup>a</sup> (8 categories) | 25.5 (15.6) | 42.3 (19.3) | 113.9 (12.3) |
| *Other cognitive domains* |  |  |  |
| Dot counting<sup>b</sup> /10 | 9.3 (4.1) | 10.0 (0) | 9.9 (0.2) |
| Position discrimination<sup>b</sup> /20 | 17.4 (3.0) | 19.5 (1.4) | 19.6 (0.9) |
| Number location<sup>b</sup> /10 | 8.8 (3.9) | 9.5 (0.9) | 9.4 (1.1) |
| Coloured progressive matrices<sup>c</sup> /36 | 19.5 (6.8) | 33.2 (5.4) | NA |
| Digit span forwards | 4.7 (1.2) | 7.2 (1.1) | 6.8 (0.9) |
| Digit span backwards | 1.8 (0.8) | 5.4 (1.1) | 4.7 (1.2) |

*These data were originally reported by Hoffman et al. (2011b). Standard deviations are shown in parentheses. <sup>a</sup> Cambridge semantic battery (Bozeat et al., 2000). <sup>b</sup> Visual Object and Space Perception battery (Warrington & James, 1991). <sup>c</sup> Raven (1982). Normative data were taken from published test norms. NA= not available.*

The two patient groups were well-matched for overall performance on the synonym judgement task (see Table 2). An SD performance index was computed for each trial by calculating the proportion of SD patients who responded correctly to the trial. This gave a measure of how difficult each trial in the test was for individuals with impaired semantic representations. The SA performance index was computed in the same way and provided us with an indicator of how difficult was for individuals with deficits in semantic control.^1^ These indices were then entered into mixed effects models as predictors of accuracy and RT in healthy participants. In other words, the SD and SA performance indices were treated as properties of trials in the task (just as frequency, imageability etc. were treated as properties of trials in the first set of analyses) and their effects on healthy adult performance were tested. Separate models were estimated for each age group as well as combined models that included effects of age group and its interactions with the other predictors.

Following the established interpretation of semantic deficits in these two groups, we assumed that a strong effect of the SD performance index would indicate that participants had weakness in the representation of semantic knowledge, whereas a strong effect of the SA performance index would indicate weakness in semantic control processes. As an additional test of these effects, we identified a subset of 20 trials in which SD patients scored substantially more poorly than SA patients (SDworst) and a subset of 21 trials in which SA patients scored substantially more poorly than SD patients (SAworst). These subsets were defined by computing a difference score for each trial (SD proportion correct minus SA proportion correct) and selecting trials where the difference exceeded 0.15 in either direction. This cut-off was chosen so that approximately 20% of trials fell into each subset. Performance of the healthy groups on these trials was then analysed.

Mixed effects models were constructed and tested using the recommendations of Barr et al. (2013). Linear models were specified for analyses of RT and logistic models for accuracy. We specified a maximal random effects structure for all models, including random intercepts for participants and items as well as random slopes for all predictors that varied within-participant or within-item. The following control predictors were included in RT models: trial order, position of target (left, centre, right), accuracy on previous trial (as errors typically lead to a pronounced slowing on the subsequent trial). Continuous predictors were standardised prior to entry in the model. The significance of effects was assessed by comparing the full model with a reduced model that was identical in every respect except for the exclusion of the effect of interest. Likelihood-ratio tests were used to determine whether the inclusion of the effect of interest significantly improved the fit of the model.

## Results

### General cognitive assessments

Two older participants scored less than 26/30 on the MMSE. Their data were excluded from the study. Scores on the background tests for the remaining participants are shown in Table 1. Older people scored slightly higher than young people on the MMSE. Young people outperformed older people on the Trail-making test, particularly part B which probes task-switching executive ability. The older group also demonstrated poorer executive function on the Wisconsin card-sorting test. In contrast, older people displayed significantly higher scores on the two tests of vocabulary, suggesting that they have greater reserves of semantic knowledge. There were no group differences on the verbal fluency tasks.

### 96-item synonym judgement test

Older people produced significantly more correct responses on the synonym judgement task (M = 97.0%; SD = 2.1) than young people (M = 89.5%; SD = 5.9; Mann-Whitney *U* = 41.5, *p* < 0.001). However, the young group were significantly faster to respond (M = 2044ms; SD = 515) compared with the older group (M = 2715ms; SD = 959; Mann-Whitney *U* = 174, *p* = 0.003). This is likely to reflect general age-related reductions in processing speed (Salthouse, 1996), rather than a semantic-specific effect. Our main interest in the present study was whether different factors influenced performance in the two age groups. Figure 2 provides the first indication that this may be the case. Accuracy on each of the 96 trials is plotted for the two groups. In both groups, the majority of trials were completed correctly by over 80% of participants. However, there were a small number of trials for which each group was less likely to give a correct response. Importantly, each group tended to fail on a different set of trials, as shown in Figure 2. Young people displayed notably poorer performance on trials that probed the meaning of low frequency words like *impetus* and *morass,* perhaps suggesting that they have yet to learn the meanings of these rarely-encountered words. In contrast, the older group performed most poorly on common words like *window* and *cause.* It is inconceivable that older people are unfamiliar with these words; instead it is likely that some aspect of the structure of these trials, such as competition between response options, which causes older people to find them particularly challenging. We test these hypotheses next.

**Figure 2:**
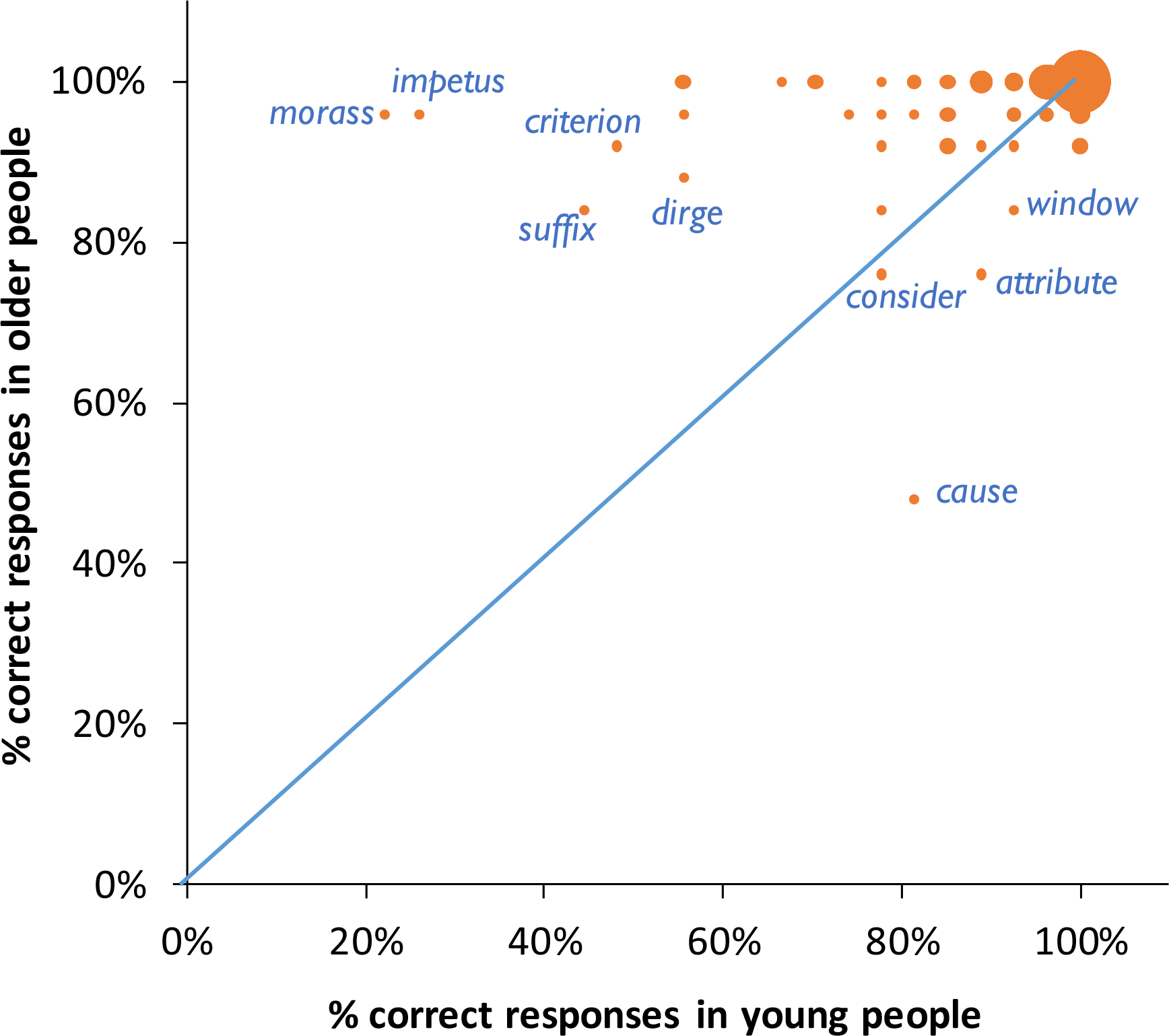
Relationship between young and older accuracy for individual trials on the test The size of the circles indicates the number of trials occupying each point. Trails falling below the diagonal were more likely to be completed correctly by young people, while trials above the diagonal were correct more often in older people.

### Using psycholinguistic variables to predict performance in young and older individuals

Correlations between all performance measures and predictors (across items) are shown in Table 3 (Spearman’s rank correlations were used due to potential ceiling effects in older adults’ accuracy; for scatterplots showing the relationship between young and older accuracy and the other predictors, see Supplementary Figure 1). Accuracy in young and older people was positively correlated, but only moderately so (*ρ* = 0.44). This suggests that, as already noted, different factors may be influencing performance in each group. Indeed, the pattern of correlations between the performance measures and the other predictors sometimes differs between the two groups. There were, for example, strong correlations between word frequency and young people’s accuracy and RT, but no such correlations for older people, and older people’s performance was correlated with semantic diversity but young people’s was not.

**Table 3:** Descriptive statistics and Spearman’s rank correlations for all dependent measures and predictors

|  | Mean (s.d.) | 1 | 2 | 3 | 4 | 5 | 6 | 7 | 8 | 9 |
| --- | --- | --- | --- | --- | --- | --- | --- | --- | --- | --- |
| 1. Young accuracy | 0.89 (0.16) | -- |  |  |  |  |  |  |  |  |
| 2. Older accuracy | 0.97 (0.07) | .44*** | -- |  |  |  |  |  |  |  |
| 3. Young RT (ms) | 2135 (680) | -.71*** | -.47*** | -- |  |  |  |  |  |  |
| 4. Older RT (ms) | 2785 (938) | -.53*** | -.56*** | .66*** | -- |  |  |  |  |  |
| 5. SD performance index | 0.72 (0.23) | .62*** | .31** | -.59*** | -.36*** | -- |  |  |  |  |
| 6. SA performance index | 0.72 (0.21) | .54*** | .39*** | -.55*** | -.50*** | .39*** | -- |  |  |  |
| 7. Probe frequency | 1.32 (0.75) | .36*** | .00 | -.39*** | -.01 | .56*** | .03 | -- |  |  |
| 8. Imageability | 450 (143) | .63*** | .43*** | .65*** | -.57*** | .38*** | .54*** | .08 | -- |  |
| 9. Semantic diversity | 1.65 (0.34) | -.07 | -.32** | .11 | .26* | .11 | -.40*** | .56*** | -.42*** | -- |
| 10. TDS | 0.34 (0.17) | .12 | .26* | -.29** | -.37*** | -.05 | .15 | -.17 | .23* | -.22* |

\* = $p < 0.05$ ; \*\* = $p < 0.01$ ; \*\*\* = $p < 0.001$ .

Parameter estimates for the mixed effects models for accuracy are shown in Table 4. The modelled effects of each predictor are plotted in the top row of Figure 3. In the combined analysis of both groups, there were strong positive effects of frequency and imageability: overall, participants tended to produce more correct responses on trials involving high frequency and highly imageable words. Importantly, however, the effect of imageability interacted with group, with young people showing a larger influence of this variable. The group x frequency interaction did not reach statistical significance (*p* = 0.07), although only the young group showed a strong effect of this variable. In the combined analysis, the main effects of semantic diversity and TDS were somewhat weaker. However, the effect of TDS interacted with group. Older people were more strongly influenced by this variable, showing a greater tendency to produce fewer correct responses on trials with low TDS values (i.e., when the target relationship was weak relative to distractor relationships). This variable did not influence the performance of young people.

**Figure 3:**
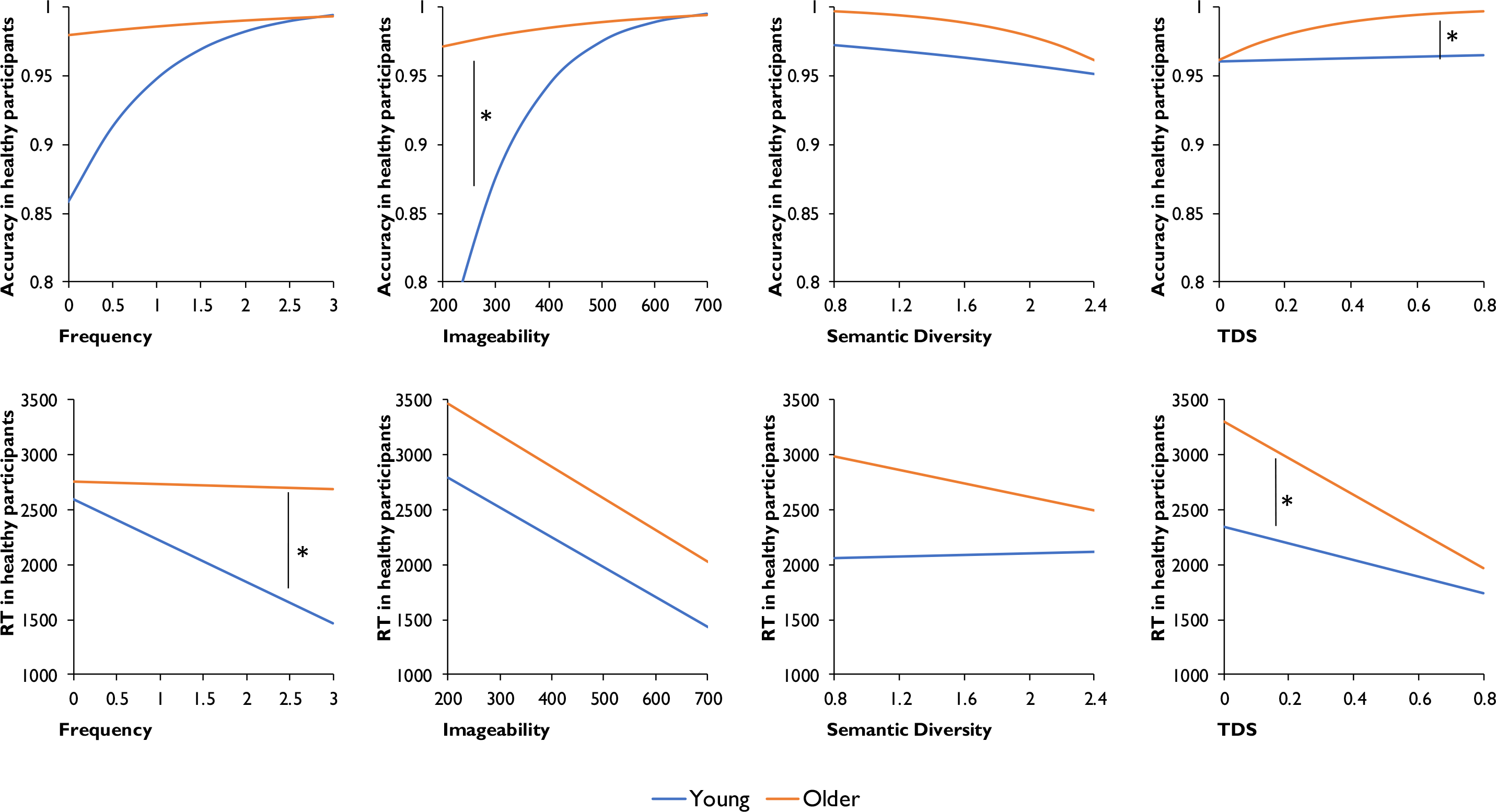
Modelled effects of psycholinguistic variables on accuracy and RT. * indicates a significant interaction between variable and group (p < 0.05).

**Table 4:**
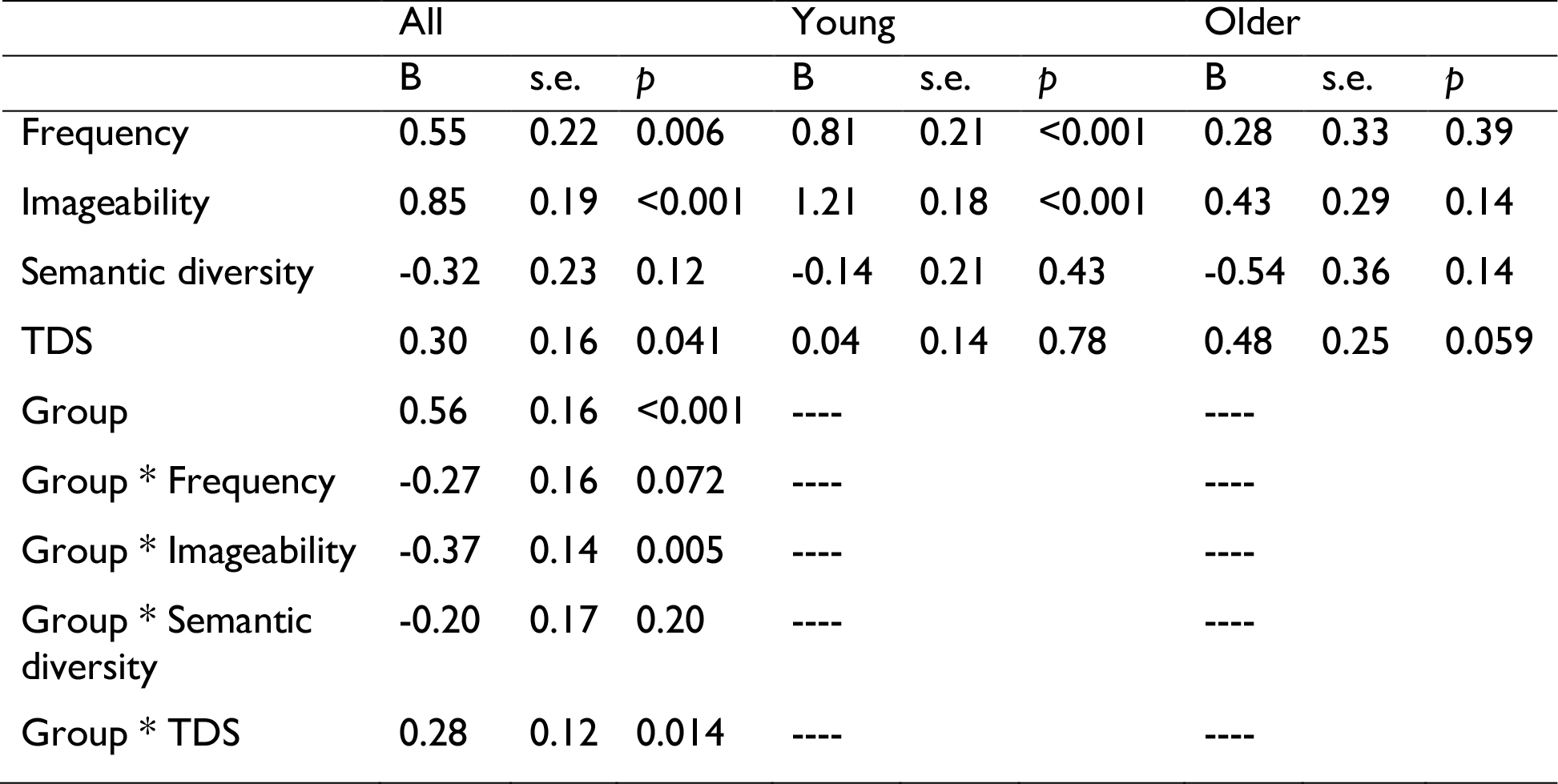
Logistic mixed effects models predicting healthy participant accuracy from psycholinguistic variables

|  | All |  |  | Young |  |  | Older |  |  |
| --- | --- | --- | --- | --- | --- | --- | --- | --- | --- |
|  | B | s.e. | p | B | s.e. | p | B | s.e. | p |
| Frequency | 0.55 | 0.22 | 0.006 | 0.81 | 0.21 | <0.001 | 0.28 | 0.33 | 0.39 |
| Imageability | 0.85 | 0.19 | <0.001 | 1.21 | 0.18 | <0.001 | 0.43 | 0.29 | 0.14 |
| Semantic diversity | -0.32 | 0.23 | 0.12 | -0.14 | 0.21 | 0.43 | -0.54 | 0.36 | 0.14 |
| TDS | 0.30 | 0.16 | 0.041 | 0.04 | 0.14 | 0.78 | 0.48 | 0.25 | 0.059 |
| Group | 0.56 | 0.16 | <0.001 | ---- |  |  | ---- |  |  |
| Group * Frequency | -0.27 | 0.16 | 0.072 | ---- |  |  | ---- |  |  |
| Group * Imageability | -0.37 | 0.14 | 0.005 | ---- |  |  | ---- |  |  |
| Group * Semantic diversity | -0.20 | 0.17 | 0.20 | ---- |  |  | ---- |  |  |
| Group * TDS | 0.28 | 0.12 | 0.014 | ---- |  |  | ---- |  |  |

Parameter estimates for the RT models are presented in Table 5, with the modelled effects of the predictors shown in the bottom row of Figure 3. There was significant effects of frequency and imageability on RT. However, only young people were slower to respond to low frequency words, with older people showing no sign of such an effect. In contrast, both groups showed similar effects of imageability, responding more slowly to less concrete words. There was a strong effect of TDS in both groups, though this effect was significantly larger in the older people. Semantic diversity had no effect on RTs.

**Table 5:** Linear mixed effects models predicting healthy participant RT from psycholinguistic variables

|  | All |  |  | Young |  |  | Older |  |  |
| --- | --- | --- | --- | --- | --- | --- | --- | --- | --- |
|  | B | s.e. | p | B | s.e. | p | B | s.e. | p |
| Frequency | -149 | 67.1 | 0.028 | -278 | 60.1 | <0.001 | -16.7 | 98.9 | 0.87 |
| Imageability | -394 | 64.9 | <0.001 | -373 | 60.1 | <0.001 | -409 | 96.9 | <0.001 |
| Semantic diversity | -44.6 | 72.7 | 0.54 | 11.0 | 64.9 | 0.87 | -102 | 106 | 0.34 |
| TDS | -207 | 51.0 | <0.001 | -126 | 44.0 | 0.005 | -288 | 76.8 | <0.001 |
| Group | 319 | 107 | 0.004 | ---- |  |  | ---- |  |  |
| Group * Frequency | 149 | 45.2 | 0.004 | ---- |  |  | ---- |  |  |
| Group * Imageability | 10.7 | 47.9 | 0.82 | ---- |  |  | ---- |  |  |
| Group * Semantic diversity | 56.3 | 49.1 | 0.25 | ---- |  |  | ---- |  |  |
| Group * TDS | 78.0 | 34.7 | 0.026 | ---- |  |  | ---- |  |  |

In summary, then, the performance of young people was most influenced by word frequency and imageability, two factors that have been linked with representational richness. In contrast, older people were more affected by the relative strength of target over distracting semantic relationships (i.e., TDS), in both accuracy and RT. These results suggest weakness in controlled semantic processing in this group. These patterns were investigated further in the next section.

### Using performance of SD and SA patients to predict performance in young and older individuals

Here, performance indices for SD and SA patients were calculated on each trial of the test (i.e., percentage of patients responding correctly to the trial) and these were used as predictors of performance in healthy individuals. Parameter estimates for the accuracy models are shown in Table 6 and the modelled effects of the predictors are presented in Figure 4. In the combined model, the SD and SA indices were strong predictors of accuracy in healthy individuals. However, the effect of the SD index interacted with group. As shown in Figure 4, young people showed a strong effect of this predictor, performing more poorly on the trials that few SD patients answered correctly. In contrast, this variable had little effect on the older group. Both groups showed similar effects of the SA index, producing fewer correct responses on trials that SA patients frequently failed.

**Figure 4:**
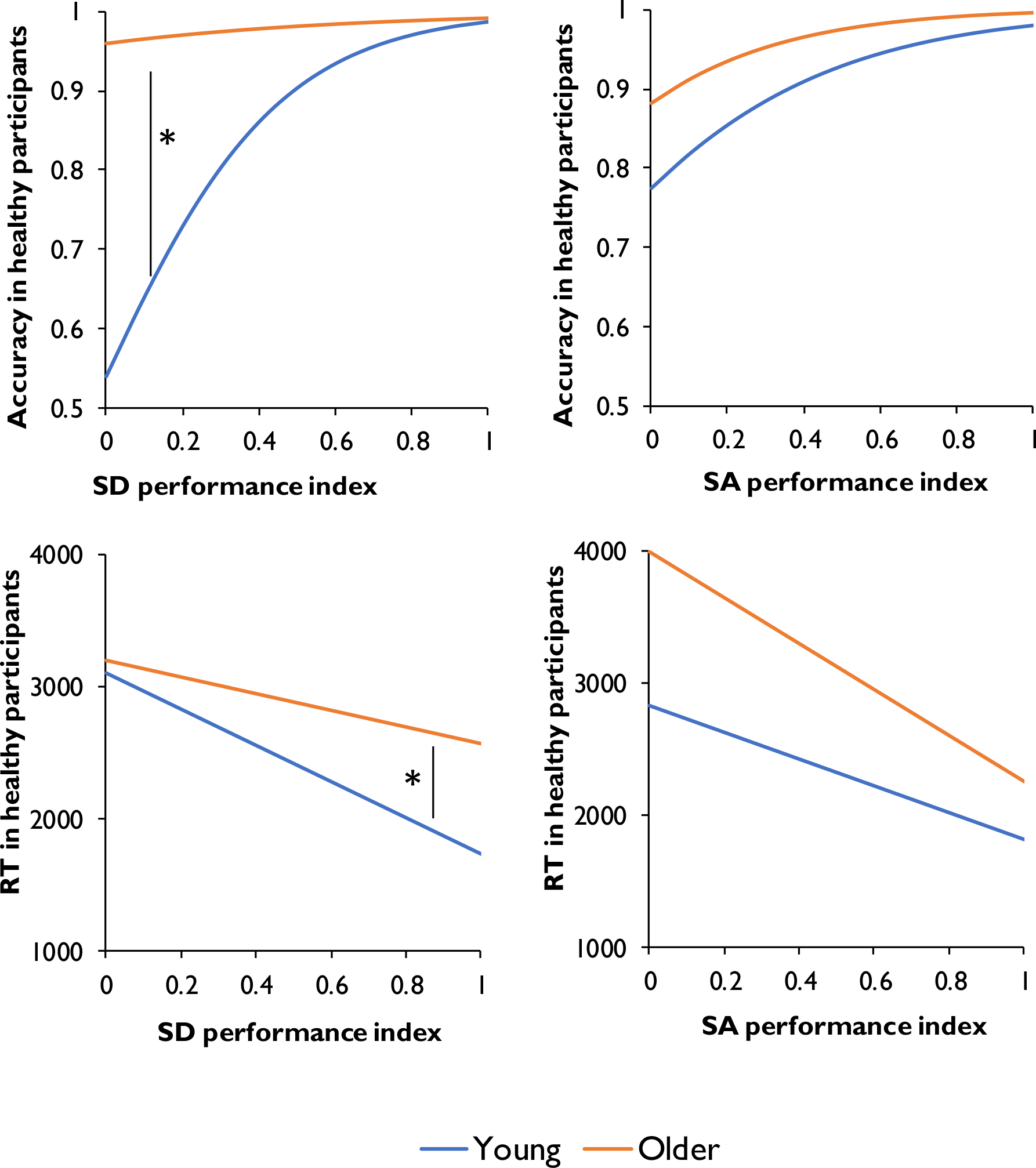
Modelled effects of patient performance indices on accuracy and RT. * indicates a significant interaction between variable and group (p < 0.05).

**Table 6:**
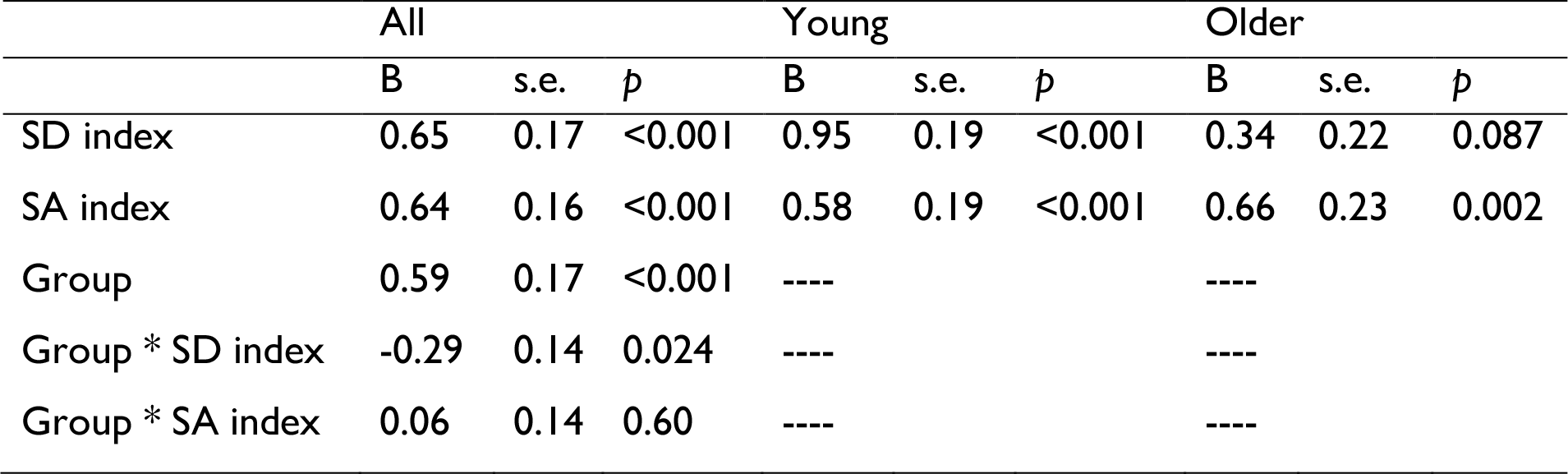
Logistic mixed effects models predicting healthy participant accuracy from patient performance indices

|  | All |  |  | Young |  |  | Older |  |  |
| --- | --- | --- | --- | --- | --- | --- | --- | --- | --- |
|  | B | s.e. | *p* | B | s.e. | *p* | B | s.e. | *p* |
| SD index | 0.65 | 0.17 | <0.001 | 0.95 | 0.19 | <0.001 | 0.34 | 0.22 | 0.087 |
| SA index | 0.64 | 0.16 | <0.001 | 0.58 | 0.19 | <0.001 | 0.66 | 0.23 | 0.002 |
| Group | 0.59 | 0.17 | <0.001 | ---- |  |  | ---- |  |  |
| Group * SD index | -0.29 | 0.14 | 0.024 | ---- |  |  | ---- |  |  |
| Group * SA index | 0.06 | 0.14 | 0.60 | ---- |  |  | ---- |  |  |

The parameter estimates for models predicting RT data are shown in Table 7. Again, SD and SA performance indices were significant predictors, with healthy participants taking longer to respond on the trials which the patient groups found most difficult. The effect of the SD performance index again interacted with group, with a stronger effect in young people (see Figure 4). The interaction of group with SA index was not statistically significant (*p* = 0.087) though there was a trend in the expected direction, i.e., a tendency for older people to show a larger effect of this predictor.

**Table 7:**
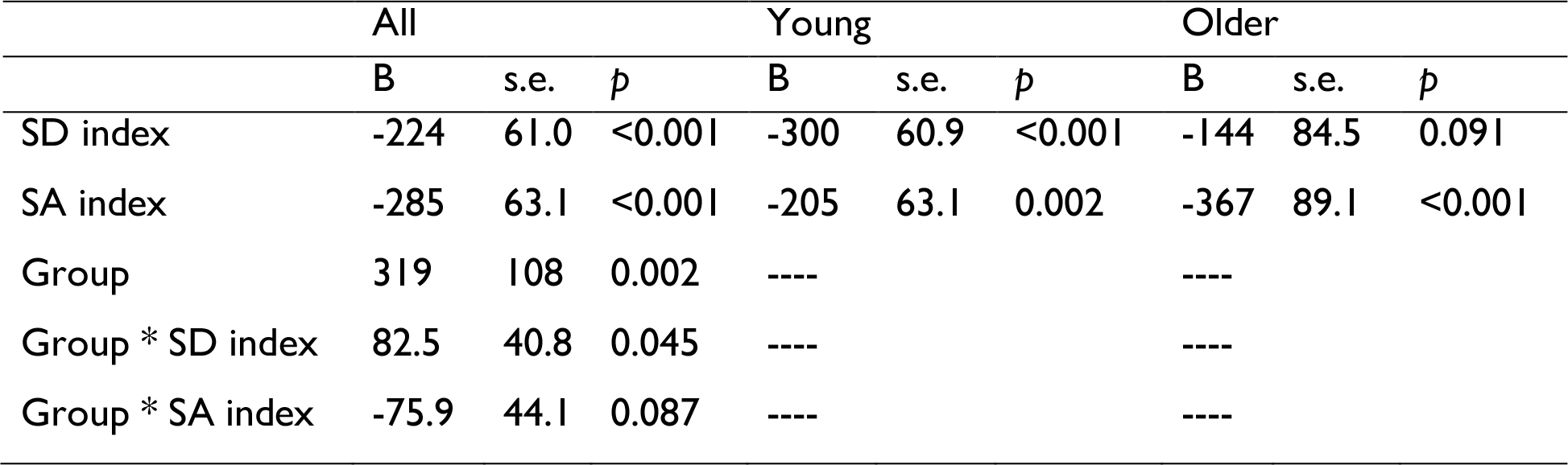
Linear mixed effects models predicting healthy participant RT from patient performance indices

To investigate these effects further, we analysed performance on subsets of trials on which one group of patients performed substantially better than the other. The results are shown in Figure 5. Young people were least accurate on the trials which SD patients found disproportionately difficult (SDworst), while older people produced fewer on the trials which SA patients found more difficult (SAworst). This difference was supported by a significant interaction between group and trial subset in a 2 x 2 mixed effect analysis (*χ*^2^ = 6.83, *p* = 0.009). For RT, older people were significantly slower to respond to the SAworst trials, compared with the SDworst trials, while there were no difference between these trial sets in young people. Again, the difference between the groups was supported by a significant 2 x 2 interaction in a mixed effects analysis (*χ*^2^ = 4.75, *p* = 0.029). These results suggest that older people’s performance was better predicted by the performance of SA patients with impaired semantic control processes, whereas the responses of young people were better predicted by the performance SD patients with impoverished semantic representations.

**Figure 5:**
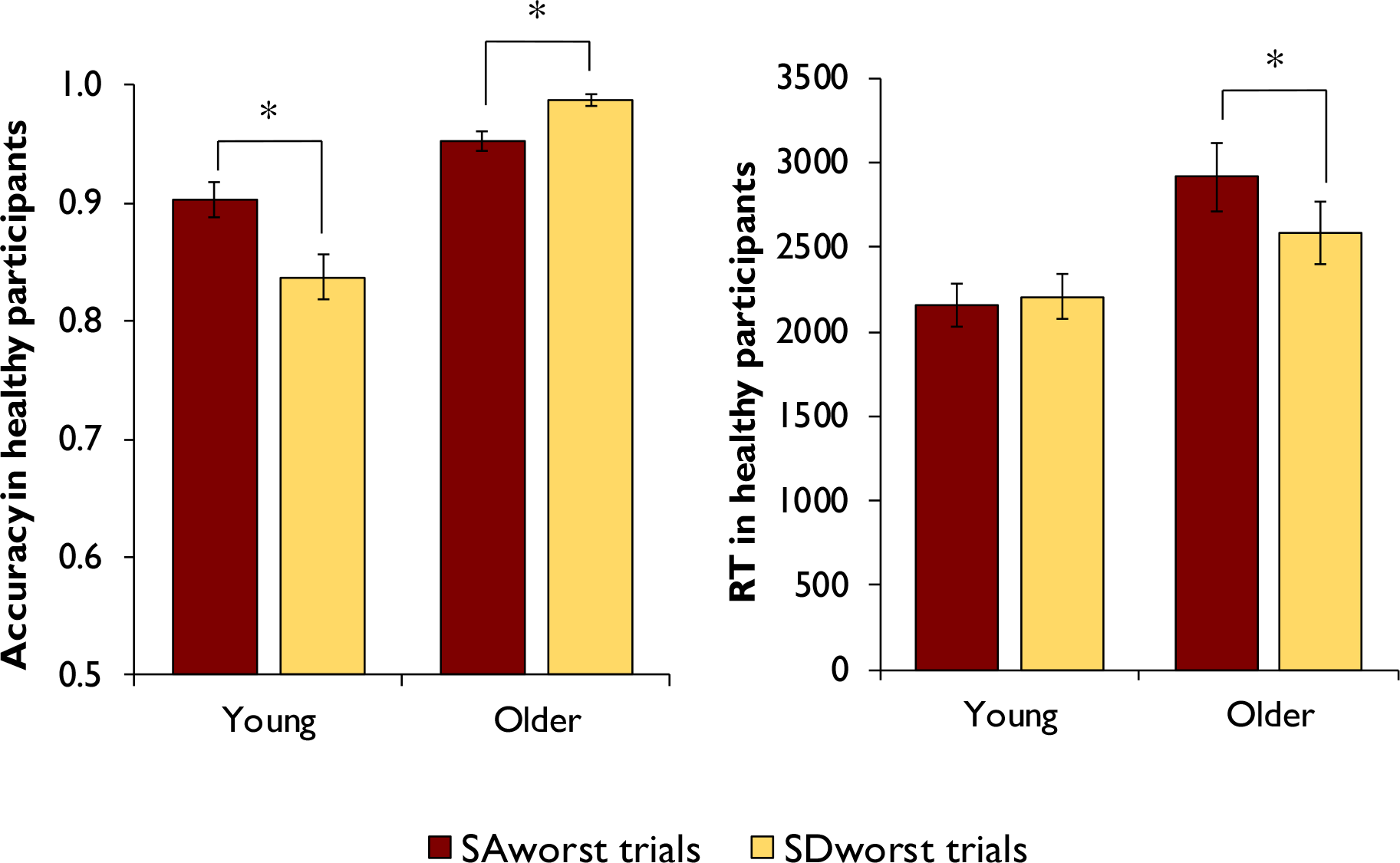
Performance on young and older people on trials for which SD or SA patients performed particularly poorly. * indicates significant within-group differences (Wilcoxon signed ranks test; p < 0.05).

## Discussion

Effective semantic cognition relies on a store of knowledge representations as well as on control processes that regulate goal-directed retrieval and manipulation of this information. We investigated the status of these capacities in young and older adults by identifying the factors that influenced their performance on a synonym-matching verbal comprehension task. Young people had difficulty processing the meanings of low frequency and abstract words, whereas these factors had less influence on the performance of older people. This indicates that the young group had smaller and less detailed repositories of semantic knowledge, as suggested by a number of previous studies (Grady, 2012; Nilsson, 2003; Nyberg et al., 1996; Park et al., 2002; Rönnlund et al., 2005; Salthouse, 2004). In contrast, older people were strongly influenced by the balance of target vs. distractor semantic relationships (TDS), performing poorly on trials where the target semantic relationship was weak relative to irrelevant relationships present in the trial. This factor had less effect in the young group, suggesting that young people are more effective at engaging semantic control processes to identify task-relevant semantic information. This picture was supported by a second analysis, in which the performance of older people was best predicted by the scores of SA patients, who had established semantic control deficits. Conversely, the performance of the young group was better predicted by that of SD patients suffering from deterioration in knowledge representations. Taken together, these findings suggest a more nuanced picture of semantic cognition in later life than has been typically assumed in the past, with more detailed knowledge representations offset by weakness in controlled processing of this knowledge. We consider the implications for each age group in turn.

The performance of young people was best predicted by the SD performance index and, like SD patients, young people had particular difficulty on trials that probed the meanings of low frequency and abstract words. What is the interpretation of these findings? The suggestion is not, of course, that young people are suffering from some sort of neurodegeneration akin to that observed in SD. Rather, the most likely explanation is that their semantic representations are still in the process of development, and the consequences of this resemble the disease-based weakening of semantic knowledge in SD. Indeed, it has often been observed that the trajectory of knowledge deterioration in SD is a mirror image of the acquisition of semantic knowledge in early development (Rogers & McClelland, 2004) and the present findings are an example of this phenomenon.

In SD, the meanings of low frequency words are thought to be particularly vulnerable to the disease process because they are represented more weakly in the knowledge store to begin with (Plaut, McClelland, Seidenberg, & Patterson, 1996; Rogers & McClelland, 2004). The semantic system has less opportunity to develop robust representations of the meanings of low frequency words because, by definition, it is exposed to them less often. This also explains why understanding of low frequency words tends to be acquired later in life (Kuperman, Stadthagen-Gonzalez, & Brysbaert, 2012; Morrison, Chappell, & Ellis, 1997; Stadthagen-Gonzalez & Davis, 2006). These same factors mean that the young adults in the present study (aged between 18 and 22) were yet to develop strong semantic representations for lower frequency words and hence showed strong effects of this variable. It is important, however, to note that young people showed strong effects of frequency in RT as well as accuracy (whereas older people showed neither). In other words, even when participants had sufficient knowledge of low frequency words to provide a correct response, they were still slower to do so. This suggests that young people’s semantic representations for low frequency words, where they do exist, are less developed than those of older people. This result is also a mirror image of semantic degradation in SD, in which concepts are not lost in all-or-nothing fashion but gradually lose detail and acuity over time (Rogers et al., 2004). It is also consistent with a recent study reported by Pexman and Yap (2018), in which participants with lower vocabulary scores showed larger frequency effects on RT when making semantic decisions.

Imageability also had a larger effect on the accuracy of young people, with this group, but not the older participants, showing poorer knowledge of more abstract words. These results also align with the pattern seen in SD. The status of imageability effects in SD is a contentious issue, with some researchers claiming that patients have *more* intact knowledge of abstract words (Bonner et al., 2009; Cousins, York, Bauer, & Grossman, 2016; Yi, Moore, & Grossman, 2007). However, on the test used in the present study, SD patients reliably show large decrements in knowledge for more abstract words (Hoffman, 2016; Hoffman & Lambon Ralph, 2011; Jefferies et al., 2009). The most likely explanation for these findings is that more concrete words develop richer semantic representations because they are associated with a wide array of sensory-motor information (Paivio, 1986). This richer representation affords them some protection from the deterioration of knowledge in SD (Hoffman et al., 2018). It is likely that this sensory richness also affects the ease of acquiring semantic representations during development (Kuperman et al., 2012; Morrison et al., 1997; Stadthagen-Gonzalez & Davis, 2006). As a consequence, young adults have less detailed representations of the meanings of abstract words.

In contrast to the findings already discussed, older people showed no effect of word frequency and weaker effects of imageability. These results suggest, in line with much previous work (Grady, 2012; Nilsson, 2003; Nyberg et al., 1996; Park et al., 2002; Rönnlund et al., 2005; Salthouse, 2004), that older people have broader and more detailed repositories of semantic knowledge, and consequently these factors have less influence on their performance. The SD performance index was not a significant predictor of performance in this group, suggesting that healthy ageing does not involve any loss of semantic knowledge of the kind seen in SD. This conclusion is consistent with data on the neuroanatomical correlates of SD and the effects of healthy ageing on cortical volumes. Knowledge loss in SD is strongly linked to atrophy of the ventral anterior temporal cortex (Butler et al., 2009; Mion et al., 2010). However, this region exhibits little age-related volume loss in healthy individuals, compared with other areas such as prefrontal cortex (Fjell et al., 2009). Indeed, a recent study in 556 healthy older adults found that volume of the ventral temporal cortices was a positive predictor of an individual’s quantity of semantic knowledge (Hoffman et al., 2017). However, this association was entirely mediated by educational level and childhood IQ, suggesting that this was a lifelong association rather than an effect of the ageing process.

The present study has, however, identified another factor that has a greater influence on semantic processing in old age. Older people showed much stronger effects of the target vs. distractor strength (TDS), a measure of the strength of the target semantic relationship on each trial relative to irrelevant distractor relationships. Older people were slower and less accurate to respond when the target semantic relationship was weak and competition from irrelevant semantic relationships was strong. Identifying the correct response under these conditions places high demands on semantic control processes that regulate the activation of semantic knowledge (Badre & Wagner, 2007; Lambon Ralph et al., 2017; Thompson-Schill, 2003). These results therefore indicate that older people are less able to exercise semantic control. In line with this view, performance in this group was strongly predicted by the SA performance index. Patients with SA have well-established deficits in semantic control (Corbett et al., 2009; Jefferies & Lambon Ralph, 2006; Noonan et al., 2010). SA patients are also strongly influenced by the semantic diversity of words, performing poorly for words that are used in a wide variety of contexts (Hoffman et al., 2011b). We observed no effect of this variable in our older adults (nor our young people) and the reason for this discrepancy is not clear, particularly as previous studies have revealed semantic diversity effects in healthy adults’ semantic processing (Hoffman & Woollams, 2015; Pexman & Yap, 2018). It may be the multiple choice format of the present task constrains the possible interpretations of the probe word’s meaning, thus mitigating the difficulty of words with highly variable meanings.

Although an old-age deficit in semantic control has not been reported previously, it is consistent with our understanding of changes in neural structure and function in later life. Semantic control is underpinned by a network of regions including inferior parietal, posterior temporal and, most prominently, inferior prefrontal cortex (Jefferies, 2013; Noonan et al., 2013; Vatansever et al., 2017). Inferior prefrontal cortex shows large age-related declines in volume in later life (Fjell et al., 2009; Raz et al., 1997; Raz et al., 2005) and changes in functional activation are also frequently observed in this region (Cabeza, 2002; Grady, 2012). Indeed, a recent neuroimaging meta-analysis indicates that older people show reduced activation during semantic processing in inferior prefrontal cortex, as well as other regions associated with semantic control (Hoffman & Morcom, 2018). These findings suggest a possible neural mechanism for the behavioural effects observed in the present study. Of course, much more work would be needed to link these behavioural and neural observations more directly and to delineate the precise circumstances under which reductions in prefrontal activity have measurable impacts on semantic performance. Potential interactions between knowledge and control should also be investigated. The age-related increases in effects of target vs. distractor strength observed in the present study suggest that conditions of high semantic competition are an area of particular weakness in later life. However, it may be that these effects are exacerbated by older people’s more expansive repositories of knowledge. It is possible that individuals who know more words, and more about those words, experience higher levels of competition when attempting to make semantic decisions. In other words, the acquisition of greater knowledge may go hand in hand with increased challenges in regulating how that knowledge is retrieved.

Finally, we note that accuracy in this study was generally high and thus our effect sizes on this measure could be limited by ceiling effects. We addressed this potential issue by using Spearman’s rank correlations to assess correlations between variables and by analysing accuracy within logistic mixed effect models (which assume a binomial rather than normal distribution of responses; see Jaeger, 2008). Nevertheless, it is possible that larger effects on accuracy would be observed in a more difficult task that elicited more errors.

## Acknowledgements

PH is supported by The University of Edinburgh Centre for Cognitive Ageing and Cognitive Epidemiology, part of the cross council Lifelong Health and Wellbeing Initiative (MR/K026992/1). Funding from the Biotechnology and Biological Sciences Research Council (BBSRC) and Medical Research Council(MRC) is gratefully acknowledged. I am grateful to Emily Hardy, Wing Yee Ho, Eszter Kalapos and Adam Ryde for assistance with data collection. Versions of this manuscript have been made available prior to publication on a preprint server (bioRxiv).

## Footnotes

1 Note that these indices were based on accuracy only: RT data were not collected from the patients.

