## Supplementary Materials for "Divergent effects of healthy ageing on semantic knowledge and control: Evidence from novel comparisons with semantically-impaired patients"

*Supplementary Figure 1: Correlations between accuracy on trials of the task and the psycholinguistic predictors and patient performance indices*

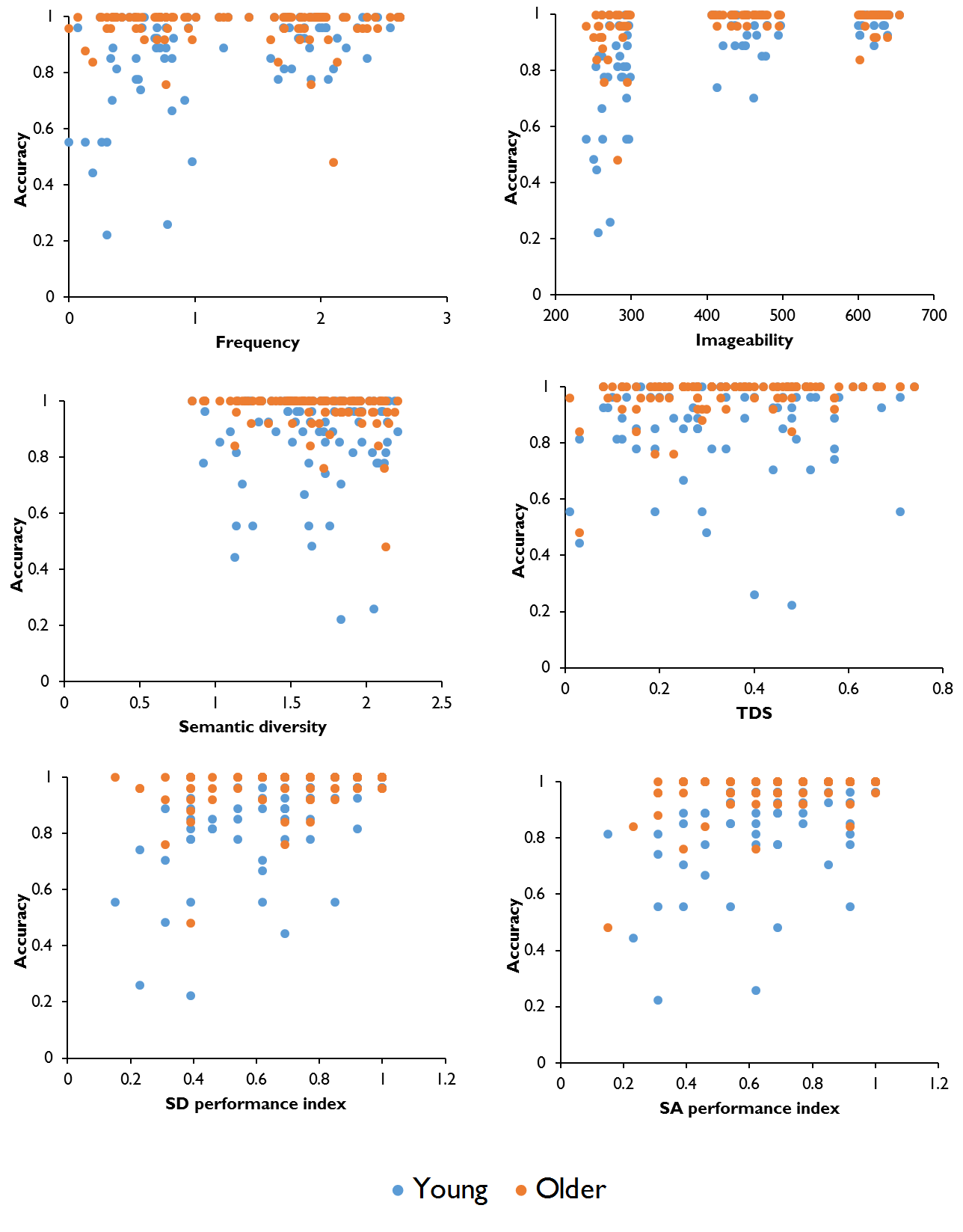
